# Clusters of fatty acids in the serum triacylglyceride fraction associate with the disorders of type 2 diabetes

**DOI:** 10.1101/279703

**Authors:** Luke W. Johnston, Zhen Liu, Ravi Retnakaran, Bernard Zinman, Adria Giacca, Stewart B. Harris, Richard P. Bazinet, Anthony J. Hanley

## Abstract

**Aims:** Our aim was to examine longitudinal associations of triacylglyceride fatty acid (TGFA) composition with insulin sensitivity (IS) and beta-cell function.

**Methods:** Adults at-risk for T2D (n=477) had glucose and insulin measured from a glucose challenge at 3 time points over 6 years. The outcome variables Matsuda index (ISI), HOMA2-%S, Insulinogenic Index over HOMA-IR (IGI/IR), and Insulin Secretion-Sensitivity Index-2 (ISSI-2) were computed from the glucose challenge. Gas chromatography quantified TGFA composition from the baseline. We used adjusted generalized estimating equations (GEE) models and partial least squares (PLS) regression for the analysis.

**Results:** In adjusted GEE models, four TGFA (14:0, 16:0, 14:1n-7, 16:1n-7 as mol%) had strong negative associations with IS while others (e.g. 18:1n-7, 18:1n-9, 20:2n-6, 20:5n-3) had strong positive associations. Few associations were seen for beta-cell function, except for 16:0, 18:1n-7, and 20:2n-6. PLS analysis indicated four TGFA (14:0, 16:0, 14:1n-7, 16:1n-7) that clustered together and strongly related with lower IS. These four TGFA also correlated highly (r>0.4) with clinically measured TG.

**Conclusions:** We found that higher proportions of a cluster of four TGFA strongly related with lower IS as well as hypertriglyceridemia, suggesting only a few fatty acids within the TGFA composition may primarily explain lipids’ role in glucose dysregulation.

## Introduction

Hypertriglyceridemia is a well described metabolic disorder resulting in negative health outcomes (1). It is a risk factor for cardiovascular disease (2, 3) and is associated with other metabolic disorders such as nonalcoholic fatty liver disease (4), the metabolic syndrome (5), and abdominal obesity (6). Circulating triacylglceride (TG) concentration is commonly measured during routine clinical assessment using enzymatic methods. However, clinically measured TG is limited as it represents the full fatty acid spectrum within the TG fraction as a summary measure. There is increasing appreciation for the importance of specific fatty acid composition profiles in different plasma fractions on various health outcomes (7–10), however there are relatively few studies that have explored the impact of the fatty acid composition in the TG fraction (11, 12).

The interaction between TG and insulin sensitivity is complex and involves components of a feedback system (3). Greater resistance to insulin in both the liver and muscle may result in greater production of TG and secretion of lipoproteins that transport TG (13). Likewise, greater TG may contribute to metabolic dysfunction and lipotoxicity in various tissues, affecting insulin sensitivity, and thus continue the cycle (3). Given the complexity and temporal nature of the relationship, long term studies with multiple data collection time points are paramount to better understanding the underlying biology and subsequent risk.

While several studies have documented prospective associations of hypertriglyceridemia with incident type 2 diabetes (T2D) (2, 14, 15), only a limited number of longitudinal studies (11, 12) have examined the relationship between TG and its composition with the pathophysiological factors underlying T2D, particularly beta-cell function. Our objective was to examine the longitudinal role of the specific composition of the serum TG fraction on OGTT-derived measures of insulin sensitivity and beta-cell function compared to clinically measured TG in a Canadian population at risk for T2D.

## Materials and Methods

Recruitment for the baseline visit of the Prospective Metabolism and Islet Cell Evaluation (PROMISE) cohort took place between 2004-2006 in London and Toronto, Canada. Individuals were selected to participate if they met the eligibility criteria of having one or more risk factors for T2D, including obesity, hypertension, family history of diabetes, and/or a history of gestational diabetes or birth of a macrosomic infant. A total of 736 individuals attended the baseline visit. Subsequent examinations occurred every three years, with data from three examination visits available for the present analysis (2004-2006, 2007-2009, and 2010-2013). The current study used data on participants who did not have T2D at baseline, who returned for one or more of the follow-up examinations, and who had samples available for fatty acid measurements (n=477; see the CONSORT diagram in Supplemental Figure S1). Metabolic characterization, anthropometric measurements, and questionnaires on lifestyle and sociodemographics were administered at each examination visit. Research ethics approval was obtained from Mount Sinai Hospital and the University of Western Ontario, and all participants provided written informed consent. Data collection methods were standardized across the 2 centres and research nurses were centrally trained.

### Metabolic characterization

After 8-12 hours of overnight fasting, participants completed a 75g oral glucose tolerance test (OGTT) at each examination visit, with blood samples taken at fasting, 30 min, and 2 hr post-glucose load. Samples were subsequently processed and frozen at −70°C. Alanine aminotransferase (ALT) was measured using standard laboratory procedures. Cholesterol, HDL, and clinically-measured TG were measured using Roche Modular’s enzymatic colorimetric tests (Mississauga, ON). Both insulin and glucose were measured from OGTT blood samples at fasting, 30 minute, and 2 hour time points. Specific insulin was measured with the Elecsys 1010 (Roche Diagnostics, Basel, Switzerland) immunoassay analyzer and electrochemiluminescence immunoassay, which shows 0.05% cross-reactivity to intact human pro-insulin and the Des 31,32 circulating split form (Linco Res. Inc) and has a coefficient of variation (CV) of 9.3%. Glucose was determined using an enzymatic hexokinase method (Roche Modular, Roche Diagnostics) with a detection range of 0.11 to 41.6 mmol/L and an inter-assay CV of <1.1% and an intra-assay CV of < 1.9%. All assays were performed at the Banting and Best Diabetes Centre Core Lab at Mt Sinai Hospital.

TGFA composition was quantified using stored fasting serum samples from the baseline visit, which had been frozen at −70°C for 4-6 years and had not been exposed to any freeze-thaw cycles. Serum fatty acids have been documented to be stable at these temperatures for up to 10 years (16). A known amount of triheptadecanoin (17:0; Nu-Chek Prep, Inc Elysian, MN, USA) was added as an internal standard prior to extracting total lipids according to the method of Folch et al (17). Each serum lipid fraction (non-esterified fatty acids (NEFA), cholesteryl ester, phospholipid, and TG) was isolated using thin layer chromatography. Fatty acid methyl esters were separated and quantified using a Varian-430 gas chromatograph (Varian, Lake Forest, CA, USA) equipped with a Varian Factor Four capillary column and a flame ionization detector. Fatty acid concentrations (nmol/ml) were calculated by proportional comparison of gas chromatography peak areas to that of the internal standards (18). There were 22 fatty acids measured in the TGFA fraction. Findings for other lipid fractions in this cohort are reported separately (see (9) for the phospholipid and cholesteryl ester fraction and (10) for the NEFA fraction analysis).

### Anthropometrics and sociodemographics

Height, weight, and waist circumference (WC) were measured at all clinic examinations using standard procedures. WC was measured at the natural waist, defined as the narrowest part of the torso between the umbilicus and the xiphoid process. BMI was calculated by dividing weight (kg) by height (m) squared. Questionnaires administered at each examination determined sociodemographics. A version of the Modifiable Activity Questionnaire (MAQ) (19) determined estimated physical activity. The MAQ collects information on leisure and occupational activity, including intensity, frequency, and duration, over the past year. Each reported activity from the MAQ was weighted by its metabolic intensity allowing for the estimation of metabolic equivalents of tasks (MET) hours per week (19).

### Variable calculation and statistical analysis

Insulin sensitivity and beta-cell function indices were computed using the OGTT glucose and insulin data. Insulin sensitivity was assessed using the Insulin Sensitivity Index (ISI) (20) and HOMA2-%S (21) using the HOMA2 Calculator. HOMA largely reflects hepatic insulin sensitivity, while ISI reflects whole-body insulin sensitivity (22). Beta-cell function was assessed using the Insulinogenic Index (23) over HOMA-IR (24) (IGI/IR) and the Insulin Secretion-Sensitivity Index-2 (ISSI-2) (25). IGI/IR is a measure of the early phase of insulin secretion while ISSI-2 is analogous to the disposition index (but is calculated using OGTT values). Each index has been validated against gold standard measures (20, 24–26).

The primary outcome variables for this analysis were HOMA2-%S, ISI, IGI/IR, and ISSI-2, which were log-transformed for the statistical modeling. The primary predictor variables for this analysis were 22 individual TGFA included as either mole percent (mol%) of the total fraction or as a concentration (nmol/mL). Clinically-measured TG was also included as a primary predictor to allow us to test the hypothesis that specific TGFA better predicted outcomes compared to clinical TG. Pearson correlation coefficients were computed to assess the relationships of individual TGFA with other continuous variables. Within TGFA composition correlations were also computed and subsequently analyzed using hierarchical clustering.

Generalized estimating equation (GEE) models (27) were used in the primary analysis to determine the longitudinal associations between the outcome variables and the predictor variables. The predictor variables were scaled (mean-centered and standardized). Given the longitudinal design, an auto-regressive of order 1 working correlation matrix was specified in the GEE model. Covariates to adjust for were selected based on the previous literature, from directed acyclic graph (DAG) (28) recommendations, and from quasi-likelihood information criteria. DAGs are used to identify the minimum adjustment necessary for a model by using the causal pathways to algorithmically identify potential confounding and colliding variables (see (28) for more detail about using DAGs). The DAG structures to understand potential confounding, shown in Supplemental Figure S2 and Supplemental Figure S3, were processed by the DAGitty software (29, 30) to generate the recommended adjustments. These DAG structures were developed based on hypothesized causal pathways between each variable, which were then input into the DAGitty software. The output from DAGitty was used, in conjunction with the other methods, to help inform the final model.

The final GEE model (M6; seen in Supplemental Table S1) was adjusted for years since baseline, WC, baseline age, ethnicity, sex, ALT, MET, and total NEFA. The variables TGFA, total NEFA, sex, ethnicity, and baseline age were classified as *time-independent* (held constant) as they were measured only at the baseline visit or do not change throughout the study, while the outcome variables and remaining covariates were set as *time-dependent*. After transformations, the GEE estimates are interpreted as an expected percent difference in the outcome variable for every standard deviation (SD) increase in the predictor variable given the covariates are held constant (including time). We also tested for an interaction with sex, ethnicity, or time by the predictor term for each outcome variable.

While GEE accounts for the longitudinal design of the data, this approach is limited in that it cannot analyze the inherent multivariate nature of the composition of the TGFA fraction. Therefore, to confirm the GEE results in a multivariate environment (i.e. all TGFA analyzed collectively), partial least squares regression (PLS) was used to identify the patterns of TGFA composition against insulin sensitivity and beta-cell function as outcome variables. Briefly, PLS is a technique that extracts latent structures (clusters) underlying a set of predictor variables conditional on a response variable(s) (i.e. the outcome variables). How accurately the clusters within the TGFA composition predict metabolic function is determined by using cross-validation on the PLS models.

A more detailed explanation of these statistical techniques and on the analysis process can be found in the supplemental methods for our paper in the NEFA fraction (10). All analyses were performed using R 3.4.4 (31), along with the R packages geepack 1.2.1 for GEE (32) and pls 2.6.0 for PLS. The R code and extra analyses for this manuscript is available at https://doi.org/10.6084/m9.figshare.5143438. Results were considered statistically significant at p<0.05, after adjusting for multiple testing using the Benjamini-Hochberg False Discovery Rate (33). STROBE was used as a guideline for reporting (34).

## Results

### Basic characteristics of the PROMISE cohort

Table 1 shows basic characteristics of the PROMISE cohort. The mean follow-up time was 5.6 (1.0) years, where 88.5% of participants attended all three visits. There were 349 (73.2%) females and 336 (70.4%) who reported European-ancestry, with a mean age in years of 50.0 (9.8) and a mean BMI of 31.1 (6.5) kg/m^2^. As expected from the study’s eligibility criteria, the majority of participants, n=305 (65.3%), had a family history of diabetes. Between the baseline visit and the 6-year visit in this sample, insulin sensitivity and beta-cell function measures had a significant median decline of between 14% to 21% (p<0.001 from GEE; n=357-470).

**Table 1:** Basic characteristics of PROMISE participants at each of the 3 clinic visits. Values as either median (interquartile range), mean (SD), or n (percent).

| Measure | Baseline | 3-yr | 6-yr |
| --- | --- | --- | --- |
| HOMA2-%S | 88.8 (54.2-136.7) | 76.8 (49.1-121.8) | 73.7 (49.5-110.1) |
| ISI | 13.6 (8.7-21.8) | 11.6 (6.9-19.1) | 11.7 (7.5-17.6) |
| IGI/IR | 7.1 (4.2-10.6) | 5.6 (3.6-9.8) | 5.6 (3.4-9.2) |
| ISSI-2 | 727.5 (570.0-922.5) | 611.2 (493.0-836.4) | 624.8 (470.0-813.8) |
| ALT (U/L) | 29.6 (16.0) | 28.4 (19.6) | 25.8 (16.9) |
| TG (mmol/L) | 1.5 (0.8) | 1.4 (0.8) | 1.4 (0.7) |
| Chol (mmol/L) | 5.2 (0.9) | 5.1 (1.0) | 5.1 (0.9) |
| HDL (mmol/L) | 1.4 (0.4) | 1.3 (0.4) | 1.4 (0.4) |
| TGFA (nmol/mL) | 3137.5 (1686.6) |  |  |
| NEFA (nmol/mL) | 383.1 (116.3) |  |  |
| MET | 46.1 (61.4) | 48.2 (60.4) | 43.7 (56.7) |
| Age (yrs) | 50.0 (9.8) | 53.2 (9.8) | 56.2 (9.6) |
| BMI (kg/m<sup>2</sup>) | 31.1 (6.5) | 31.4 (6.5) | 31.1 (6.6) |
| WC (cm) | 98.5 (15.5) | 99.2 (15.7) | 100.5 (15.8) |
| Ethnicity |  |  |  |
| - European | 336 (70%) |  |  |
| - Latino/a | 59 (12%) |  |  |
| - Other | 50 (10%) |  |  |
| - South Asian | 32 (7%) |  |  |
| Sex |  |  |  |
| - Female | 349 (73%) |  |  |
| - Male | 128 (27%) |  |  |

Figure 1 shows the composition of each FA in the TG fraction (see Supplemental Table S2 for the raw values). Three TGFA contributed 82.4% to the total TG concentration: 18:1 n-9 (37.8%); 16:0 (26.6%); and, 18:2 n-6 (18.0%). Figure 2 shows a heatmap of the correlation of individual TGFA as concentrations with the outcome variables and several basic characteristics. As expected, nearly all TGFA had very strong positive correlations (r= 0.34 to 0.92) with clinically-measured TG and moderate positive correlations with WC (r=0.31 to 0.36). There were also moderate negative correlations with HDL (r=−0.53 to −0.31). For the outcome variables, the correlations for the insulin sensitivity measures were generally higher (HOMA2-%S: r=-0.47 to −0.32, ISI: r=−0.47 to −0.31) than for the beta-cell function measures (all r<0.30). For correlations of individual TGFA using mol% with the basic participant characteristics, as shown in Figure 3, differences in correlations between fatty acids were most evident for 14:0, 14:1n-7, 16:0, and 16:1n-7 that had a moderate positive correlation with clinical TG (r=0.42 to 0.52) while all other fatty acids had a negative association (r=-0.5 to −0.34). In particular, those fatty acids with the negative associations with clinical TG were all the very long chain polyunsaturated fatty acids (e.g. 20:4n-6, 20:5n-3). As seen in Figure 4, four fatty acids (14:0, 16:0, 14:1n-7, and 16:1n-7) clustered together, each highly positively correlated with each other and negatively correlated with all other fatty acids.

**Figure 1:**
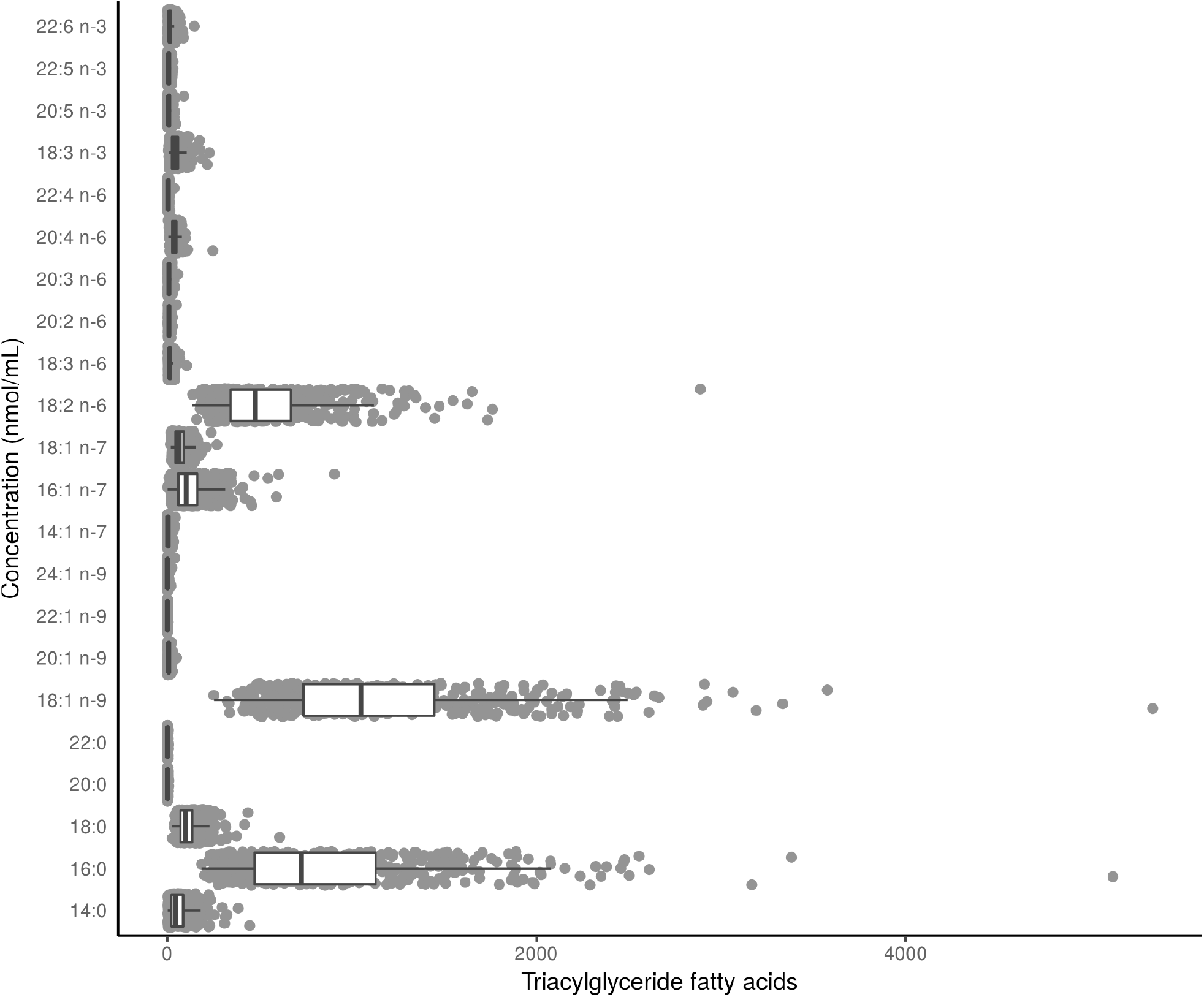
Distribution of the composition of TGFA in the baseline visit of PROMISE participants (2004-2006). Boxplots represent the median and interquartile range of the FA values.

**Figure 2:**
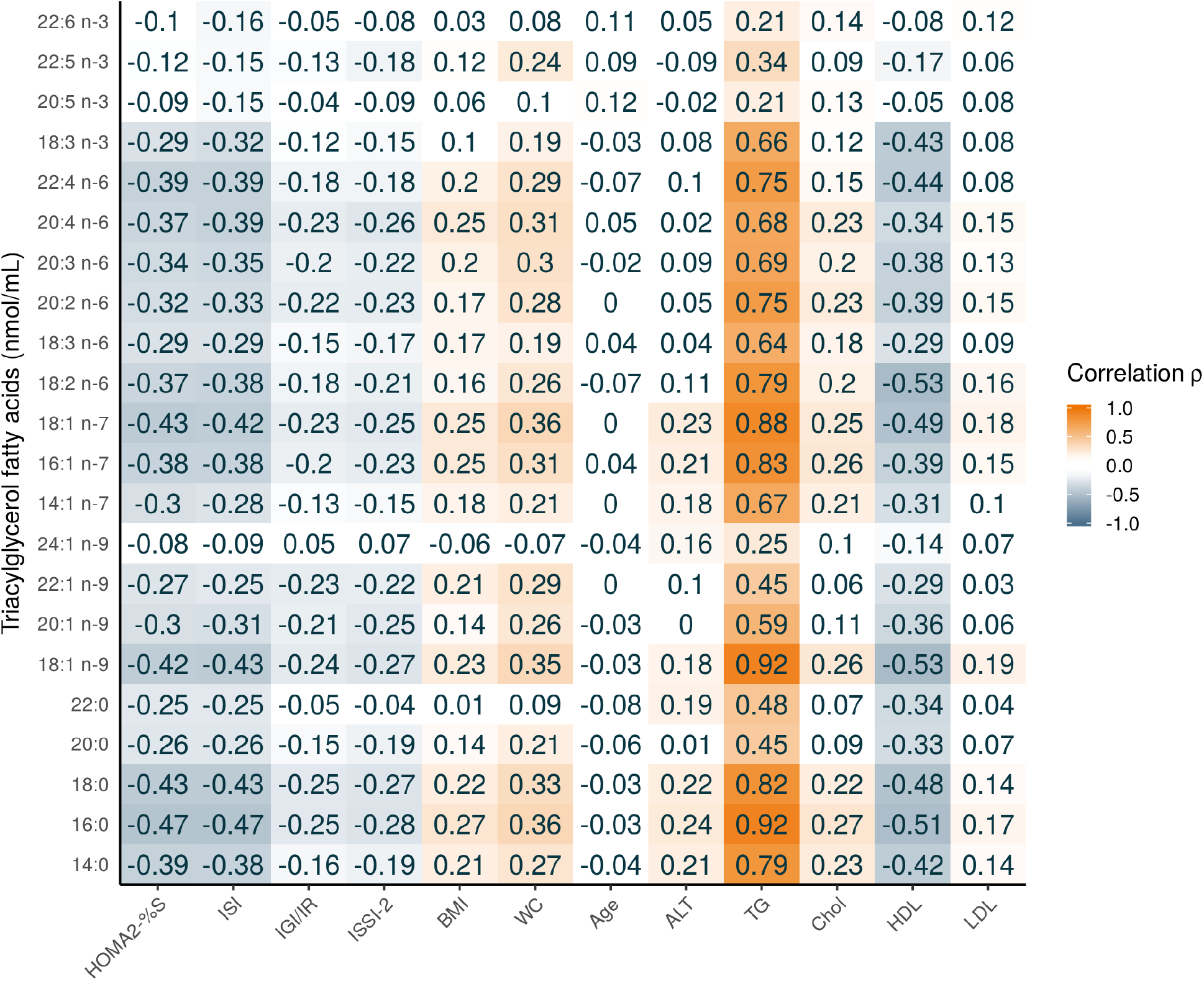
Pearson correlation heatmap of TGFA (nmol/mL) with continuous basic and metabolic characteristics of PROMISE participants from the baseline visit (2004-2006). Darker orange represents a positive correlation; darker blue represents a negative correlation.

**Figure 3:**
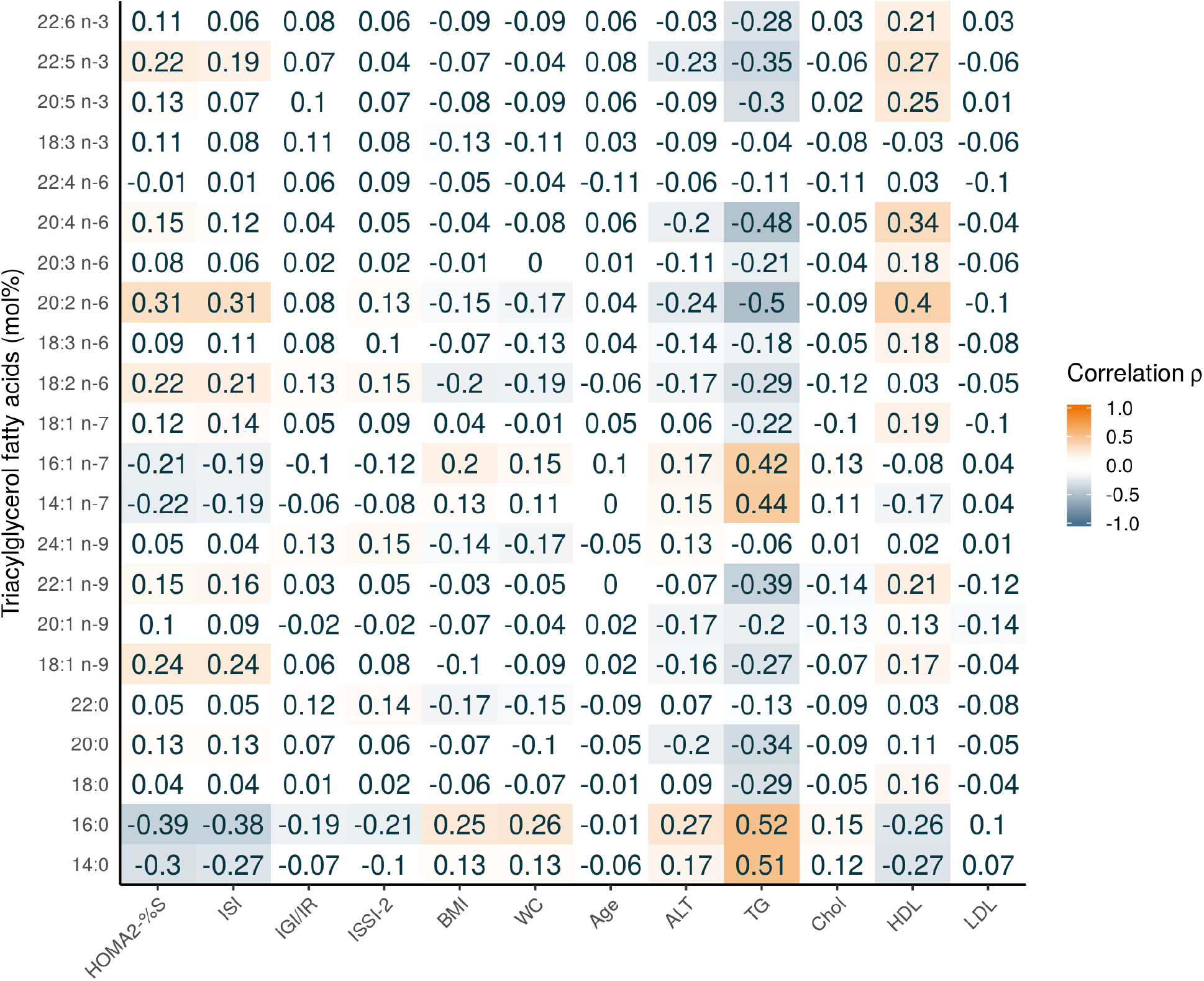
Pearson correlation heatmap of TGFA (mol%) with continuous basic and metabolic characteristics of PROMISE participants from the baseline visit (2004-2006). Darker orange represents a positive correlation; darker blue represents a negative correlation.

**Figure 4:**
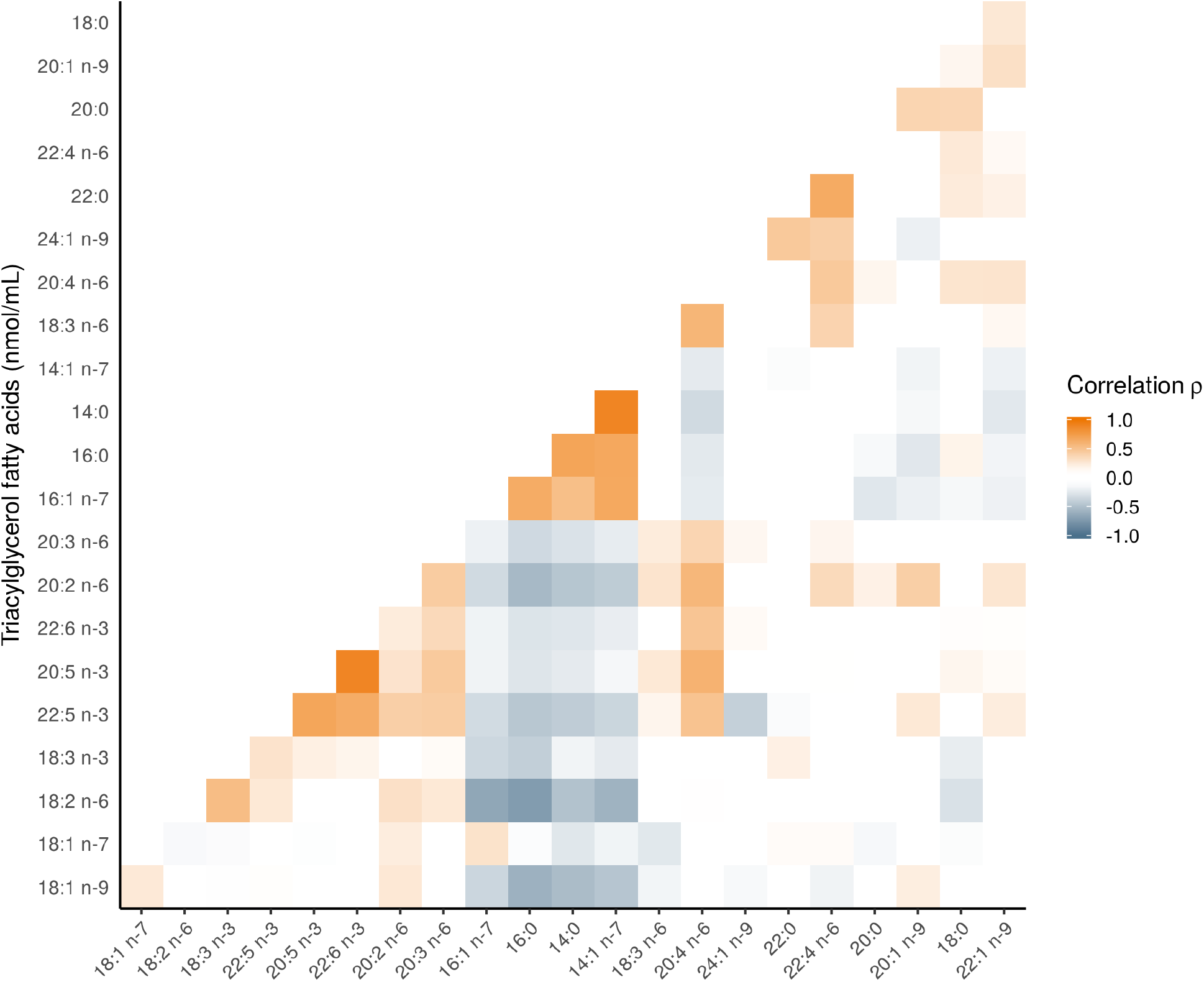
Pearson correlation heatmap of TGFA in the PROMISE participants from the baseline visit (2004-2006). The correlations of FA grouped using heirarchical cluster analysis; FA along the x and y axis are ordered according to this analysis. Darker orange represents a positive correlation; darker blue represents a negative correlation.

### Generalized estimating equation models

Results from the unadjusted GEE model are shown in Figure 5 and for the adjusted GEE model in Figure 6. The majority of associations with beta-cell function measures were attenuated after full model adjustment, while nearly all associations with insulin sensitivity remained significant for both mol% and nmol/mL results. Subsequent analysis revealed that the attenuation with beta-cell function was due primarily to adjustment for WC.

**Figure 5:**
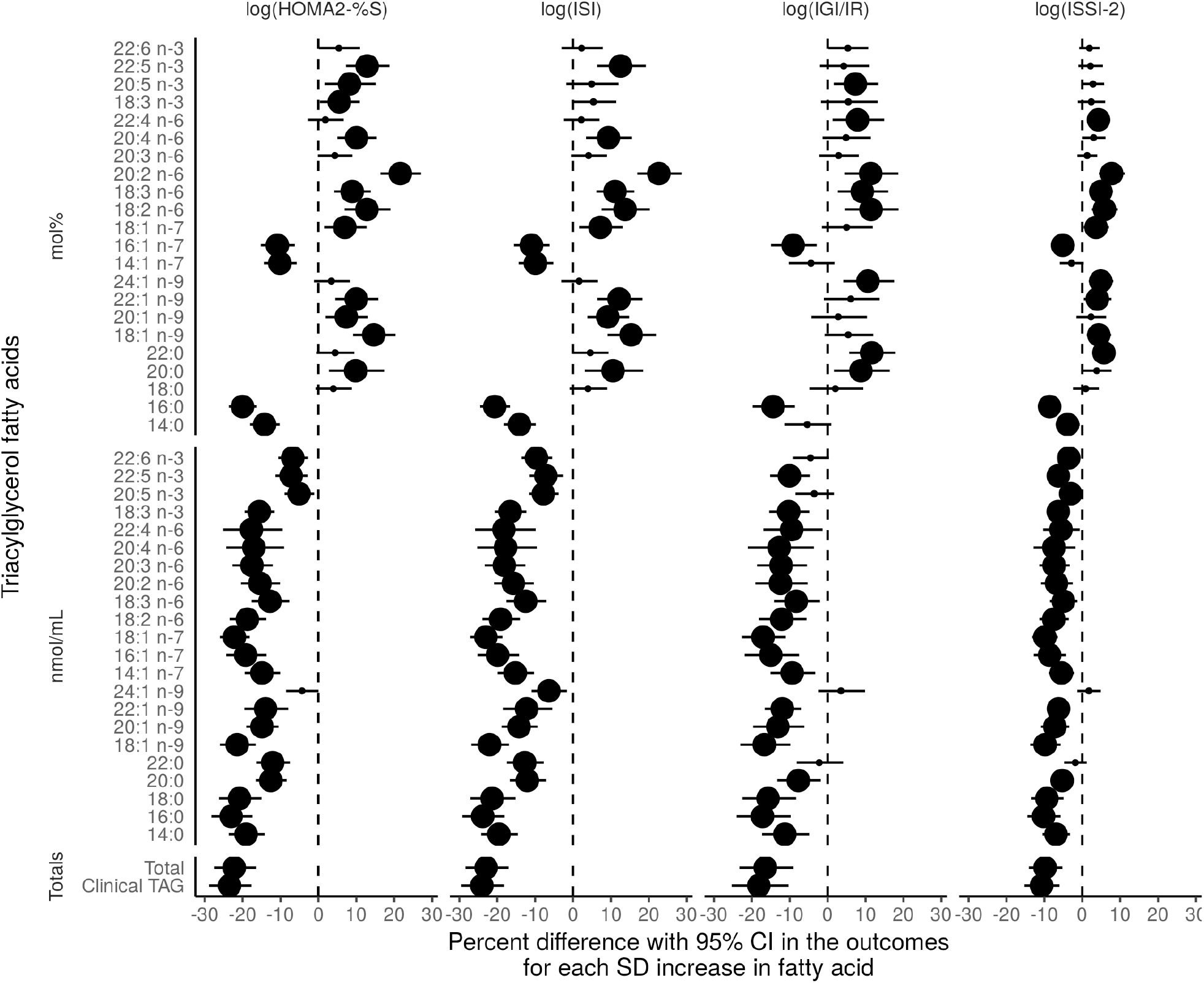
Time-adjusted GEE models of the association of the TGFA (mol% and nmol/mL) and total clinically-measured TG with insulin sensitivity and beta-cell function outcomes using the 6 year longitudinal data from the PROMISE cohort. X-axis values represent a percent difference in the outcome per SD increase in the FA. P-values were adjusted for the BH false discovery rate, with the largest dot representing a significant (p<0.05) association.

**Figure 6:**
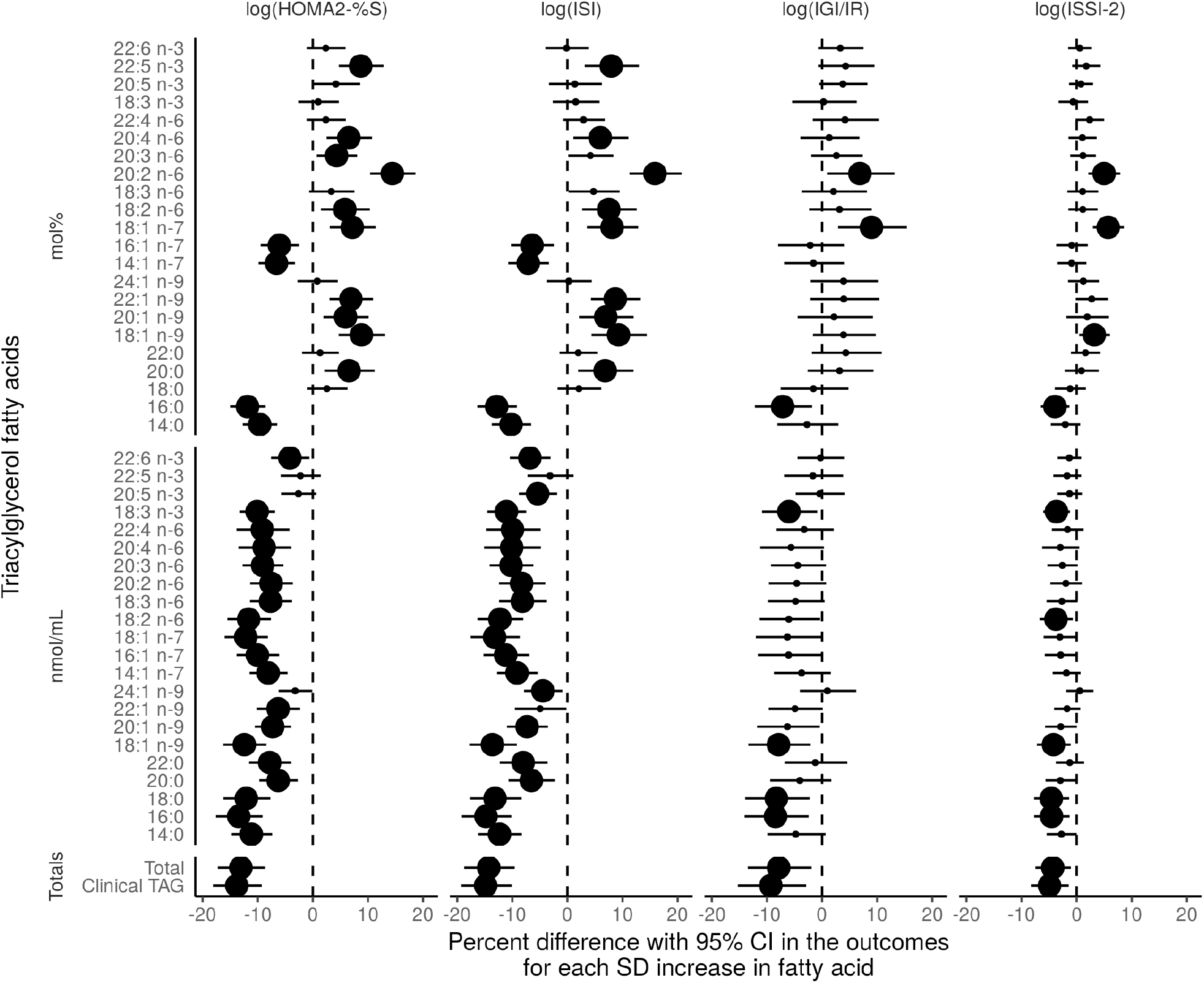
Fully-adjusted GEE models of the association of the TGFA (mol% and nmol/mL) and total clinically-measured TG with insulin sensitivity and beta-cell function outcomes using the 6 year longitudinal data from the PROMISE cohort. Variables controlled for were follow-up time, WC, baseline age, ethnicity, sex, ALT, physical activity, and total NEFA. X-axis values represent a percent difference in the outcome per SD increase in the FA. P-values were adjusted for the BH false discovery rate, with the largest dot representing a significant (p<0.05) association.

In analyses using concentration values, nearly all TGFA had a strong negative association on HOMA2-%S and ISI (estimates of percent difference ranging from −13.5 to −4.2 and −14.8 to −4.5, respectively), and a few had strong negative associations with IGI/IR and ISSI-2 (estimates ranging from −8.4 to −6.0 and −4.6 to −3.7, respectively). In analyses using TGFA mol% values, four TGFA (14:0, 16:0, 14:1n-7, and 16:1n-7) had negative associations with HOMA2-%S and ISI (estimates between −11.9 to −6.1 and −12.8 to −6.4, respectively, lower insulin sensitivity for every SD increase in the TGFA), while several more TGFA had positive associations with HOMA2-%S and ISI (20:0, 18:1n-9, 20:1n-9, 22:1n-9, 18:2n-6, 20:2n-6, 20:4n-6, and 22:5n-3) estimating between 4.3 to 14.4 and 5.9 to 15.9%, respectively, higher insulin sensitivity for every SD increase in the TGFA. One TGFA, 20:2n-6, had a very strong positive association with the insulin sensitivity measures, with a 14.4 to 9.3% higher insulin sensitivity for every SD increase. Both clinically-measured TG and total TGFA concentration had very strong negative associations with all outcome variables.

While there were a few significant interactions by time in unadjusted models, after inclusion of covariates in the model, these interactions were attenuated (data not shown). There were no significant interactions by sex or ethnicity for any of the TGFA (data not shown). Results of the sensitivity analyses identifying WC as the covariate that attenuated the beta-cell function associations from the unadjusted model are shown in Supplemental Figure S4. A tabular presentation of the GEE results is shown in Supplemental Table S3 for unadjusted models and Supplemental Table S4 for adjusted models.

### Clustering of TGFA by metabolic measures

The PLS analysis corroborated the findings from the GEE models. The PLS results conditioned on insulin sensitivity as the outcome showed a clustering of the fatty acids 14:0, 14:1n-7, 16:0, and 16:1n-7 as mol% (Figure 7). These TGFA loaded strongly and negatively on HOMA2-%S and ISI in the first component, suggesting this cluster of TGFA tracks together with lower insulin sensitivity. The TGFA 20:2n-6, 20:5n-3, 22:5n-3, and 22:6n-3 loaded positively on both insulin sensitivity measures. No other TGFA loaded strongly. In the second component, 18:1n-9 and 18:1n-7 loaded positively but not strongly while 20:5n-3 and 22:6n-3 loaded strongly and negatively with both HOMA2-%S and ISI; however, this component only explained <10% of the variance. The PLS model for insulin sensitivity had good predictive ability, with a high correlation between the predicted outcome values against the observed values (HOMA2-%S: r=0.46, p<0.001; ISI: r=0.39, p<0.001).

**Figure 7:**
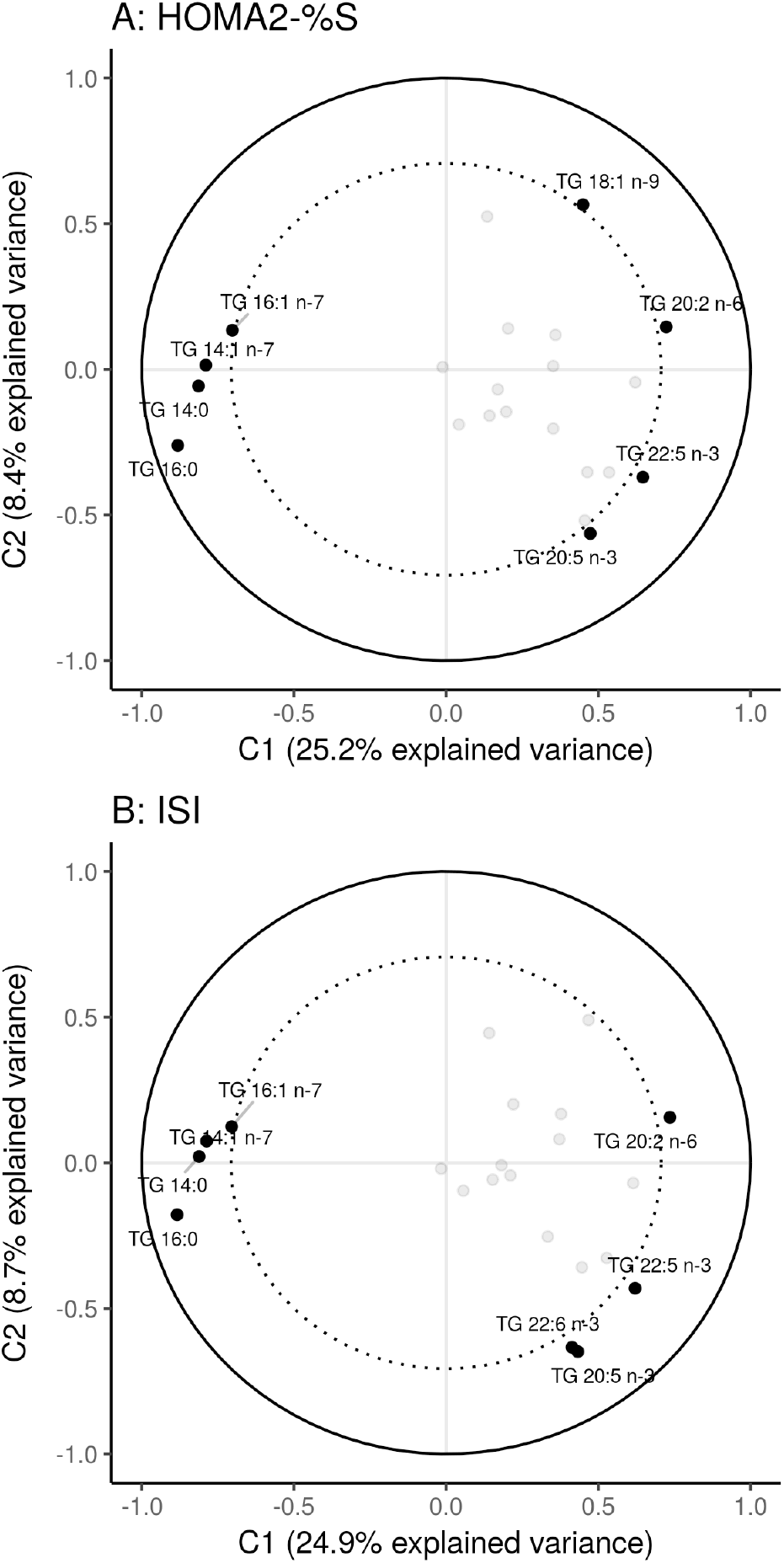
Partial least squares (PLS) models showing the clustering of TGFA on insulin sensitivity and beta-cell function measures. The percent explained variance of each component is shown in brackets on each axis. The solid line represents an explained variance of 100% while the dashed line represents an explained variance of 50%. FA between these lines represent variables that strongly explain the underlying structure of the data.

The beta-cell function PLS results showed a similar clustering of fatty acids, however there was a lower correlation (though significant at p<0.001) between the predicted values and the observed values (r=0.25 to 0.24), suggesting that TGFA composition poorly predicts beta-cell function. Given the low predictability, only the insulin sensitivity measures are presented. We used the extracted PLS scores as the predictor variable in the GEE models and found negative associations of the first component on all outcome variables, with the strongest association being with the insulin sensitivity variables (estimate of 10.4, all p<0.001; using PLS scores constrained by ISI). See Supplemental Figure S5 for a plot of the loadings of each TGFA on the two components.

## Discussion

In the present study, we found that in a Canadian cohort at risk for T2D, several specific TGFA and groups of TGFA were strongly associated with insulin sensitivity and moderately associated with beta-cell function. In particular, the TGFA myristic acid (14:0), 7-tetradecenoic acid (14:1n-7), palmitic acid (16:0), and palmitoleic acid (16:1n-7) all strongly and negatively associated with lower insulin sensitivity. While most TGFA were not associated with beta-cell function, three fatty acids, palmitic acid (16:0), *cis*-vaccenic acid (18:1n-7), and eicosadienoic acid (20:2n-6) were associated negatively and positively, respectively, with measures of beta-cell function. Using PLS, we also found that four TGFA (14:0, 14:1n-7, 16:0, 16:1n-7) clustered together, and that this cluster strongly predicted lower insulin sensitivity.

To our knowledge, no longitudinal study to date has examined the role of the composition of the TGFA fraction on detailed OGTT-derived metabolic measures. Two large prospective studies have been published that similarly examined TGFA composition and T2D outcomes. Rhee *et al* presented a nested case-control analysis (n=189 cases and n=189 controls) within the Framingham offspring cohort (11), which found that subjects with a TGFA composition characterized by a lower carbon chain and fewer double bonds (e.g. 14:0, 16:0) had a higher risk for T2D after 12-years while those with a profile characterized by higher carbon chain and more double bond TGFA had a lower risk for T2D. A similar pattern of TGFA was also associated with HOMA-IR cross-sectionally at the baseline visit. In addition, Lankinen *et al* reported on a prospective cohort of males in Finland (12), for which TGFA data were available for 831 participants after 6-years of follow-up. In their cohort, OGTT data were only available at the 6-year visit. In the cross-sectional analysis they found that most saturated fatty acids had negative associations with insulin sensitivity and beta-cell function while linoleic acid (18:2n-6), docosapentaenoic acid (22:5n-3), eicosapentaenoic acid (20:5n-3), and arachidonic acid (20:4n-6) had positive associations with insulin sensitivity. The magnitude of the associations were larger for the insulin sensitivity results compared to the beta-cell function results, similar to what we observed. Our study extends these findings by using multiple measurements of metabolic function and as well as multivariate statistical approaches that allowed us to identify clusters of TGFA. In another much smaller study (n=16) of mostly females (35), the authors reported a positive correlation between total esterified (of which TG make up the majority) 16:0, 16:1n-7, and 18:1n-9 with HOMA-IR, findings which were largely similar to the present analysis.

There are a few possible explanations for these findings. Circulating TGFA derive from three sources: adipose lipolysis, dietary fat, and de novo lipogenesis (DNL). Dietary carbohydrates and fat can influence DNL activity (36–38). Determining the specific source of TGFA is extremely difficult to ascertain outside of highly controlled experimental settings. Many previous studies that have examined DNL as the source have used markers of estimated DNL, such as the ratio between 18:2n-6 to 16:0 or 16:1n-7 to 16:0 (12, 39, 40). However, there are limitations to using these ratios as the fatty acids used in their calculation can also be obtained from the diet in addition to being created through DNL (39). An experimental feeding trial (n=24) was conducted to identify the fatty acids that most accurately reflected DNL as potential biomarkers (41). The study found that palmitoleic acid (16:1n-7), directly measured DNL using isotopes, and liver fat were all highly correlated with each other (r>0.50), suggesting that 16:1n-7 may be a good biomarker for hepatic DNL. In another small (n=14) feeding trial, meal type (high fat vs low fat) was tested to determine its effect on DNL and TGFA composition (42). The authors reported that 14:0, 16:0, 16:1, and 18:2 were higher in the low fat (high carbohydrate) group. In another recent overfeeding trial, 1000 kcal of saturated fat, unsaturated fat, or carbohydrates over 3 weeks was given to 38 overweight individuals (38) to test changes in liver fat and hepatic DNL. At the end of the study, the carbohydrate group had higher DNL activity as well as an increase in liver fat, though the saturated fat group had the highest increase in liver fat. This link between carbohydrate intake and DNL activity has been well documented (1, 3, 4, 37, 39, 43, 44).

In our findings, the four fatty acids were highly positively correlated amongst each other and negatively or neutrally with all other TGFA, in addition to clustering together on their negative association with insulin sensitivity. This may suggest that greater DNL activity is the source of these TGFA. Several studies have shown a link between higher estimated DNL activity and an increased risk for metabolic dysfunction (8, 12, 40, 45). How DNL may influence metabolic dysfunction is not well understood. Possible reasons may be that higher DNL produces more of certain fatty acids or that higher DNL increases circulating TG, which itself is well documented to contribute to metabolic dysfunction and which we found in our study as a high positive correlation between the four TGFA and clinical TG.

Regardless of the exact source of these fatty acids, our results, in addition to the available scientific evidence, emphasize the importance of the fatty acid composition on metabolic health, as individual fatty acids can have specific physiological functions. For instance, a higher concentration of circulating 14 and 16 carbon fatty acids may expose tissues to greater lipotoxicity, for instance from palmitic acid (16:0), which is well-known to have harmful effects on tissues (46, 47). Our study extends these findings by showing that TGFA with 14 to 16 carbons clustered together and this pattern strongly associated with lower insulin sensitivity. While some of these fatty acids also had a significant association with beta-cell function, the magnitude of associations were more modest compared to those for insulin sensitivity.

The direction of association between TGFA and insulin sensitivity is unclear from previous cross-sectional studies due to the physiological feedback mechanisms involved. For example, while higher TGFA may promote muscle insulin resistance, the reverse may also be true (48). As we found no interaction by time of TGFA on insulin sensitivity, this study cannot determine the exact role of the feedback mechanism. However, combining the lack of a time interaction and the consistent negative association in models without the time interaction, these results at least suggest that the feedback mechanism may not be strongly influential and that TGFA may predict insulin sensitivity at least over a six year period. Given the complex biological mechanisms and feedback loops involved, disentangling whether insulin sensitivity influences TGFA more strongly than TGFA influencing insulin sensitivity will require more complex research designs and analyses.

Given the close biological relationship between circulating NEFA and TG, NEFA may act as a confounding factor and was thus adjusted for. In our published analysis of the NEFA fraction (10), we found that higher total NEFA, but not the specific composition, associated with lower beta-cell function. This is in contrast to the TGFA findings that the specific composition does differentially associate with insulin sensitivity and beta-cell function, adjusting for total NEFA. There was no difference in results in models that did not include NEFA as a confounder (data not shown). This difference in results between NEFA and TGFA suggests that TGFA may independently and strongly influence the pathophysiology of T2D, when compared to other lipid fraction compositions, including the phospholipid and cholesteryl ester fractions (9). This may be due to TG being biologically destined for uptake by non-hepatic tissue as they are found mainly in VLDL, at least during fasting. This is in contrast to NEFA that are mostly taken up by the liver and used in TG production (49).

Our study has potential limitations that need to be considered when interpreting the results. Firstly, this is an observational cohort and as such there may be some residual confounding we were not able to control for or were unaware of. However, we have taken extensive, empirically based precautions in identifying potential confounders and mediators through the use of the DAG modeling, relying on previous literature, and through information criteria model fit comparison methods. Only fasting TGFA were quantified, and only at the baseline visit. TGFA composition can fluctuate substantially throughout the day, so in order to control for this, PROMISE participants came for the clinic visit in the morning and fasted. There is some evidence to suggest that fasting TG is better able to discriminate diabetes cases compared to a postprandial state (11). Because TGFA were only measured at the baseline visit, we cannot investigate whether there are concomitant changes in TGFA and the metabolic measures over time. However, to optimally use GEE to analyze the data and for interpretation, we used the model to infer that a given value of TGFA could predict values of insulin sensitivity or beta-cell function over a 6 year period. This in our view is a strength of our analysis, as it reduces the chance of reverse causality given the tight integration of the glucose and fatty acid metabolism pathways, as well as maximizes the specific usage of the GEE modeling.

PLS is a well-established technique for constructing predictive models of high dimensionality data structures (i.e. fatty acid composition), however a limitation is that the initial models analyzed through PLS and the final computed scores are not able to control for potential confounders and other effect modifiers. PLS is also not able to handle longitudinal data so only the baseline visit was used in the PLS analysis, although we analyzed the extracted scores using the GEE modeling to overcome this limitation and observed concordant results between the PLS and GEE analyses.

Our study has several notable strengths, including the longitudinal design and the use of advanced statistical techniques for data analysis. These statistical techniques take advantage of the longitudinal data to allow appropriate investigation of temporal relationships and are able to handle the multidimensional nature of the data. Lastly, our cohort contains highly detailed and comprehensive variable measurements for the fatty acids and outcomes, of which were collected at each visit.

## Conclusion

In conclusion, we found that a TGFA composition containing higher proportions of 14:0, 14:1n-7, 16:0, and 16:1n-7 associated strongly with lower insulin sensitivity and (more moderately) with lower beta-cell function. We also found that most other TGFA (e.g. 20:0, most omega 6 and 9 TGFA, and 20:5n-3) associated positively with insulin sensitivity. Only a few TGFA associated positively and consistently with beta-cell function (e.g. 18:1n-7, 20:2n-6). These results provide more insight into how individual TGFA contribute to the pathogenesis of T2D while reinforcing the importance and value of clinically measured total TG as an indicator of metabolic health.

## Acknowledgements/grant support

The authors thank Jan Neuman, Paula Van Nostrand, Stella Kink, Nicole Rubio, and Annette Barnie of the Leadership Sinai Centre for Diabetes, Mount Sinai Hospital, Toronto, Canada and Sheila Porter and Mauricio Marin of the Centre for Studies in Family Medicine, University of Western Ontario, London, Canada for their expert technical assistance and dedication in their work for PROMISE.

## Contribution statement

The authors had the following responsibility: LWJ conducted research, analyzed data, and wrote the paper; RR, BZ, ZL, and SBH designed research, conducted research, and provided essential materials (infrastructure and clinical resources); RR, BZ, SBH, RPB, and AG provided intellectual feedback on the paper; RPB and ZL conducted research, provided essential reagents and materials; AJH designed research, assisted with interpretation, and provided intellectual feedback on all versions of the paper; LWJ and AJH had primary responsibility for final content. All authors read and approved the final manuscript.

## Abbreviations

ALT: Alanine aminotransferase
CV: Coefficient of variation
DNL: De novo lipogenesis
DAG: Directed acyclic graph
GEE: Generalized estimating equations
HOMA2-%S: Homeostatic model of assessment 2 - percent sensitivity
IGI/IR: Insulinogenic index over homeostatic model of assessment for insulin resistance
IS: Insulin sensitivity
ISI: Insulin sensitivity index
ISSI-2: Insulin Secretion-Sensitivity Index-2
MAQ: Modified activity questionnaire
MET: Metabolic equivalent of task
NEFA: Non-esterified fatty acids
OGTT: Oral glucose tolerance test
PROMISE: Prospective Metabolism and Islet Cell Evaluation cohort
PLS: Partial least squares
SD: Standard deviation
TGFA: Triacylglycerol fatty acids
WC: Waist circumference

